# Decoupling of plastid and endomembrane homeoviscous response to low temperature and darkness in *Phaeodactylum tricornutum*

**DOI:** 10.1101/2025.04.24.650388

**Authors:** Yannick Sérès, Camille Serbutoviez-Verville, Rachel Bonnarde, Marion Schilling, Mathilde Simon, Juliette Salvaing, Juliette Jouhet, Eric Maréchal, Alberto Amato

## Abstract

Understanding how marine diatoms adapt to fluctuating environmental conditions is critical for deciphering their ecological success. Here, we investigated the lipidomic responses of the model diatom *Phaeodactylum tricornutum* to low temperatures (10 °C), darkness, and combined stress. Using a detailed lipidomics approach, we revealed significant remodeling of plastidial and extraplastidial lipid species. Plastidial lipids showed increased unsaturation levels, particularly the 16:3-to-16:4 fatty acid switch, only under light conditions. This light-dependent adaptation suggests a tightly regulated mechanism that aligns lipid remodeling with photosynthetic activity.

Phospholipids in the endomembrane system exhibited less dependency on light, with notable changes even under extended dark periods. These findings highlight distinct strategies employed by diatoms to maintain membrane fluidity and protect cellular processes during stress. Unlike plants, *P. tricornutum* lacks polygalactosyldiacylglycerols, underscoring a divergent adaptation pathway.

Our results demonstrate that *P. tricornutum* adapts to cold and dark environments through lipid composition changes, supporting survival in variable marine habitats. This study provides critical insights into diatom resilience, paving the way for understanding their role in global biogeochemical cycles.

**Highlight:** Low temperature and darkness trigger distinct lipidomic remodeling in *Phaeodactylum tricornutum*, revealing species-specific adaptive mechanisms that maintain membrane integrity and enhance resilience under extreme environmental conditions.

## Introduction

Glycerolipids provide building blocks for the bulk of most biological membranes in eukaryotes. They self-organize as bilayers, each glycerolipid being mobile laterally with more or less fluidity. They consist of a glycerol backbone, derived from glycerol-3-phosphate (G3P), with one or two fatty acids (FA) esterified at positions *sn-*1 and *sn-*2, and bearing a polar head group at position sn-3, which defines specific lipid classes. Position sn-3 may also hold a third acyl group, forming triacylglycerol, which is excluded from membranes and accumulates in long-term carbon storage organelles known as lipid- or oil droplets. Based on the carbon chain-length of acyl groups and their desaturation levels, each glycerolipid class can contain various molecular species, that are essential for their structural and functional properties (Dolch and Maréchal, 2015). In photosynthetic organisms, such as plants and algae, FA synthesis occurs in the stroma of the plastid through the action of FA synthase of type II (FASII) (Brown *et al*., 2009), producing fully saturated acyl chains reaching 14 to 16 carbon chain-lengths in algae and 16 to 18 in plants. *De novo* synthesized FA are thioesterified to a stromal acyl carrier protein (ACP). Following synthesis saturated FAs can be monounsaturated by the action of a soluble desaturase acting on acyl-ACP (for review, Dolch and Maréchal, (2015)). Saturated or monounsaturated FAs can be esterified to G3P to provide plastid lipids, such as monogalactosyldiacylglycerol (MGDG), digalactosyldiacylglycerol (DGDG), sulfoquinovosyldiacylglycerol (SQDG), and phosphatidylglycerol (PG). Alternatively, they can be released by a thioesterase and exported to the extraplastidial environment where they undergo further elongation and desaturation and integration in the structure of extraplastidial lipids.

A decrease in temperature reduces lateral molecular motion that increases membrane rigidity (Garab *et al*., 2022), affects membrane permeability (Mills and Needham, 2005; Ghysels *et al*., 2019; Frallicciardi *et al*., 2022) and can have possible negative effects on cell survival. As part of a general counteracting response, lipid remodeling can be triggered, adjusting their physical properties (reviewed by Shomo *et al*. (2024)). Indeed, when temperature decreases, the proportion of long-chain polyunsaturated FA (LC-PUFAs) is often observed in various lipid classes, making them more fluid in membranes, as part of a still poorly comprehended homeoviscous adaptation mechanism (Ernst *et al*., 2016). The significance of LC-PUFAs is widely recognized (Geneste and Faure, 2022) for their role in enhancing resilience to low temperatures (Miquel *et al*., 1993; Zheng *et al*., 2016; Wang *et al*., 2021).

In the environment, a decrease in temperature can occur within minutes or hours, particularly at the end of the day, when night falls. Phytoplankton may also encounter a decrease in temperature coinciding with long darkness exposure during the descent or sinking in the water column. Cross-talks between cell responses to temperature decrease and long darkness exposures are therefore expected. Lipid remodeling also plays a role in the response to prolonged darkness, involving distinct mechanisms. In this context, lipophagy is activated as an adaptive strategy to address the absence of carbon fixation (reviewed by Amari *et al*. (2024)).

In the ocean, the wide variability in temperature and light availability significantly influences phytoplankton physiology and adaptive mechanisms (Mann and Lazier, 2013). In the oceanic ecosystems, diatoms play essential ecological roles and are key contributors to various biogeochemical cycles of elements (Benoiston *et al*., 2017; B-Béres *et al*., 2023). Diatoms, members of the Stramenopile (or Heterokonta) clade, include a diverse group of unicellular organisms that arose from a secondary endosymbiotic event (Moustafa *et al*., 2009) approximately 180–250 million years ago (Sorhannus, 2007). This event involved the engulfment of a red algal ancestor by a heterotrophic eukaryote, with the red alga subsequently establishing itself as a permanent photosynthetic organelle within the host. Over evolutionary time, genes from the red algal endosymbiont were integrated into the nuclear genome of the host, facilitating the integration of the symbiont and resulting in highly chimeric genomes (Nisbet *et al*., 2004; Bowler *et al*., 2008). Secondary plastids in these organisms are typically surrounded by four membranes, with diatom plastids characterized by loosely packed thylakoid membranes (Flori *et al*., 2016, 2017). This intricate cellular architecture, along with their complex, composite genomes, likely contributed to the evolutionary success of diatoms, establishing them as one of the most ecologically dominant clades in marine environments (Benoiston *et al*., 2017). Diatoms colonized other moist environments, including both submerged and subaerial habitats in marine and freshwater systems. Diatoms are distributed across a broad latitudinal range (Massana *et al*., 2014; Seeleuthner *et al*., 2018), inhabiting low and high latitude waters (Thaler and Lovejoy, 2014). Given their wide distribution, diatoms have to cope with permanently changing physico-chemical contexts and have developed intriguing strategies to respond *e.g.* to nitrate (Abida et al., 2015; Rogato *et al*., 2015; Dolch et al., 2017, Busseni *et al*., 2019), phosphate (Abida et al., 2015; Alipanah *et al*., 2018; Dell’Aquila and Maier, 2020; Dell’Aquila *et al*., 2020), iron (Allen *et al*., 2008) starvations, long periods out of the euphotic zone (Nymark *et al*., 2013) due to mixing of the water column (Mellor and Durbin, 1975) and to polar nights (Morin *et al*., 2020; Joli *et al*., 2024).

The average temperature at the sea surface in temperate oceanic regions ranges from about 8 to 20 °C, based on data collected in the World Ocean Atlas (Reagan *et al*., 2023). In the present investigation we have exposed non-acclimated cultures of the model diatom *Phaeodactylum tricornutum* to four days at 10 °C (leaving the photoperiod and irradiance unvaried), or complete darkness or a combination of the two conditions and analyzed the glycerolipidome response in detail in comparison to cultures kept at 20 °C and exposed to a control photoperiod (12h:12h light:dark photoperiod and an irradiance of 40 µmol photons·m^-2^·s^-1^).

## Materials and methods

### Maintenance and culturing of Phaeodactylum tricornutum

The *P. tricornutum* strain Pt1 8.6 (CCAP 1055/1; CCMP2561 from the Culture Collection of Marine Phytoplankton, now known as NCMA: National Center for Marine Algae and Microbiota) is routinely maintained under slow growth conditions in the laboratory on solid ESAW10N10P medium with 1% m/v agar. The cultures are stored at 20 °C in a PHCbi MLR-352-PE Climate Chamber (PHC Group, Tokyo, Japan) under constant illumination at 50 µmol photons·m^-2^·s^-1^. The ESAW10N10P medium formulation follows the protocols established by Abida *et al*. (2015) and Guéguen *et al*. (2024), with a modified ESAW recipe originally described by Harrison *et al*. (1980) and Berges *et al*. (2002), where the NaNO_3_ concentration is increased tenfold. Sodium phosphate (NaH_2_PO_4_), as per Gagneux-Moreaux *et al*. (2007), is also used at a tenfold higher concentration. For experimental purposes, liquid cultures were handled under standardized procedures to ensure reproducibility. Cells were gradually transferred from solid to liquid cultures, starting with 25-mL Pyrex® glass flasks containing 5 mL of ESAW10N10P medium, followed by sequential transfers to 50-mL (10 mL), 100-mL (20 mL), and 250-mL (50 mL) flasks. Cultures were incubated in an INFORS HT Multitron Pro incubator with the following settings, temperature at 20 °C, agitation at 100 rpm, a 12:12 hour light:dark cycle, and an irradiance of 40 µmol photons·m^-2^·s^-^1. The positive control was inoculated at a density of 0.5 × 10□ cells·mL□¹ and cultured under optimal conditions (as stated above) for four days until harvest. For the cold-light condition, triplicate flasks were inoculated at 1 × 10□ cells·mL□¹ and incubated at 10 °C for four days before harvesting. For the dark exposure experiments (both cold and 20 °C conditions), triplicate flasks were inoculated at 0.5 × 10□ cells·mL□¹ and initially cultured under optimal conditions for four days. Subsequently, the cultures were transferred to darkness under the respective temperature conditions for an additional four days before harvesting.

The higher inoculum for the dark and double stress conditions is justified by the fact that, in darkness, cell division arrests compared to light conditions. Therefore, to obtain sufficient biomass for lipid extraction, a higher inoculum was needed. The low temperature was set at 10 °C and the dark was obtained by wrapping flasks into three layers of aluminum foil. After a four-day exposure to the desired condition, cells were counted and harvested then lipids were extracted and analyzed as described in the next section. The experiments were carried out three times using independent cell batches in order to take into account the natural variability of glycerolipidome analyses at the high-performance liquid chromatography-tandem mass spectrometry (HPLC-MS/MS).

### Measurement of cell density

Cell density was quantified using a TECAN Infinite® M1000 PRO (Tecan Group Ltd., Männedorf, Switzerland) spectrophotometer, following the procedure outlined by Conte *et al*. (2018). A 300 µL aliquot from each culture was transferred into a 96-well flat-bottom clear plate (ThermoFisher Scientific). Absorbance was recorded at 730 nm and converted into cell concentration using the following linear regression equation:

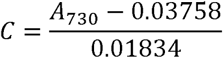

where *C* represents the cell concentration and *A*_730_ is the absorbance at 730 nm.

### Lipidomics analyses

Total glycerolipid extraction was carried out using an EDGE® automated extraction system (CEM), employing a protocol based on the method developed by Folch *et al*. (1957) with slight modifications tailored to *Phaeodactylum tricornutum* to improve efficiency. A total of 150×10□ cells were collected by centrifugation at 3,500 ×g for 9 minutes and transferred into 1.7 mL SafeSeal Microcentrifuge Tubes (27210, Sorenson). The cell pellets were boiled at 95 °C for 15 minutes to inactivate lipases. The boiled pellets were stored at −80 °C for at least one night. Prior to extraction, the samples were freeze-dried overnight using a CHRIST Alpha 2-4 LSCbasic system. One twentieth of the total glycerolipid extract was subjected to methanolysis. Lipids were dissolved in 1 mL of ACS-grade chloroform (Honeywell Riedel-de Haën), and a 50 µL aliquot was transferred to a 10-mL crimp-cap vial (Gerstel) using a Hamilton pipette. For the methanolysis reactions, two different conditions were employed: with and without heating. The former produces FA methyl-esters of all the FAs esterified to lipids and the latter produces FAMEs from free FAs. In both cases, sulfuric acid as a catalyst was used. Methanolysis was automated using a MultiPurpose Sampler (MPS, Gerstel). A fixed quantity of internal standard (5 µg of capillary-GC-grade pentadecanoic acid (15:0) (Sigma, P6125), prepared as a 1 mg·mL□¹ solution in 1:2 v/v chloroform/methanol), was introduced by the robot in each sample. The gas chromatography-flame ionization detection (GC-FID) (Perkin Elmer Clarus 580 with a 30-m cyanopropyl polysilphenesiloxane column, 0.22 mm diameter) and high-performance liquid chromatography-tandem mass spectrometry (HPLC-MS/MS) were performed according to previously established methods (Dolch *et al*., 2017; Jouhet *et al*., 2017; Guéguen *et al*., 2024).

The level of unsaturation of the different lipid classes was quantified using the double bond index (DBI). The DBI represents the average number of double bonds present in the fatty acid chains of glycerolipid molecular species. The DBI was calculated following the formula (Zheng *et al*., 2011)

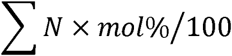

Where *N* is the total number of double bonds in the acyl chains esterified to each lipid molecular species and *mol*% is the percentage of each lipid species within each class. Multiple t-test and statistical significance determined using the Bonferroni-Dunn method, with alpha = 0.05 (without assuming a consistent standard deviation) were performed using GraphPad Prism version 7.04 for Windows, GraphPad Software, Boston, Massachusetts USA, www.graphpad.com. Adjusted p-values are reported in each figure using asterisks.

## Results

### Effects of cold and darkness on lipid classes and fatty acid profile

*Phaeodactylum tricornutum* cultures exposed to low temperature (10 °C), darkness, or a combination of both for four days exhibited a statistically significant increase in phosphatidylcholine (PC) and phosphatidylethanolamine (PE), accompanied by a corresponding decrease in galactoglycerolipids. Levels of mono- and digalactosyldiacylglycerols (MGDG and DGDG) declined under all conditions, with the exception of MGDG in the cold-dark treatment. Triacylglycerol (TAG) content was almost undetectable in both dark conditions (Fig. 1A), consistent with the observation that TAG synthesis in diatoms is predominantly driven by neosynthesis rather than lipid remodeling. Because it has been demonstrated that exposure to cold induces the accumulation of phosphatidic acid (PA) in plants, a well-established signaling molecule (Wu *et al*., 2022, and literature therein), we investigated the presence of this metabolite in our samples, despite its absence from previous reports in *P. tricornutum* (Abida *et al*., 2015). As expected, PA was not detected in the glycerolipidome of *P. tricornutum*, suggesting that PA levels in diatoms are remarkably low and/or subject to rapid turnover.

**Figure 1.**
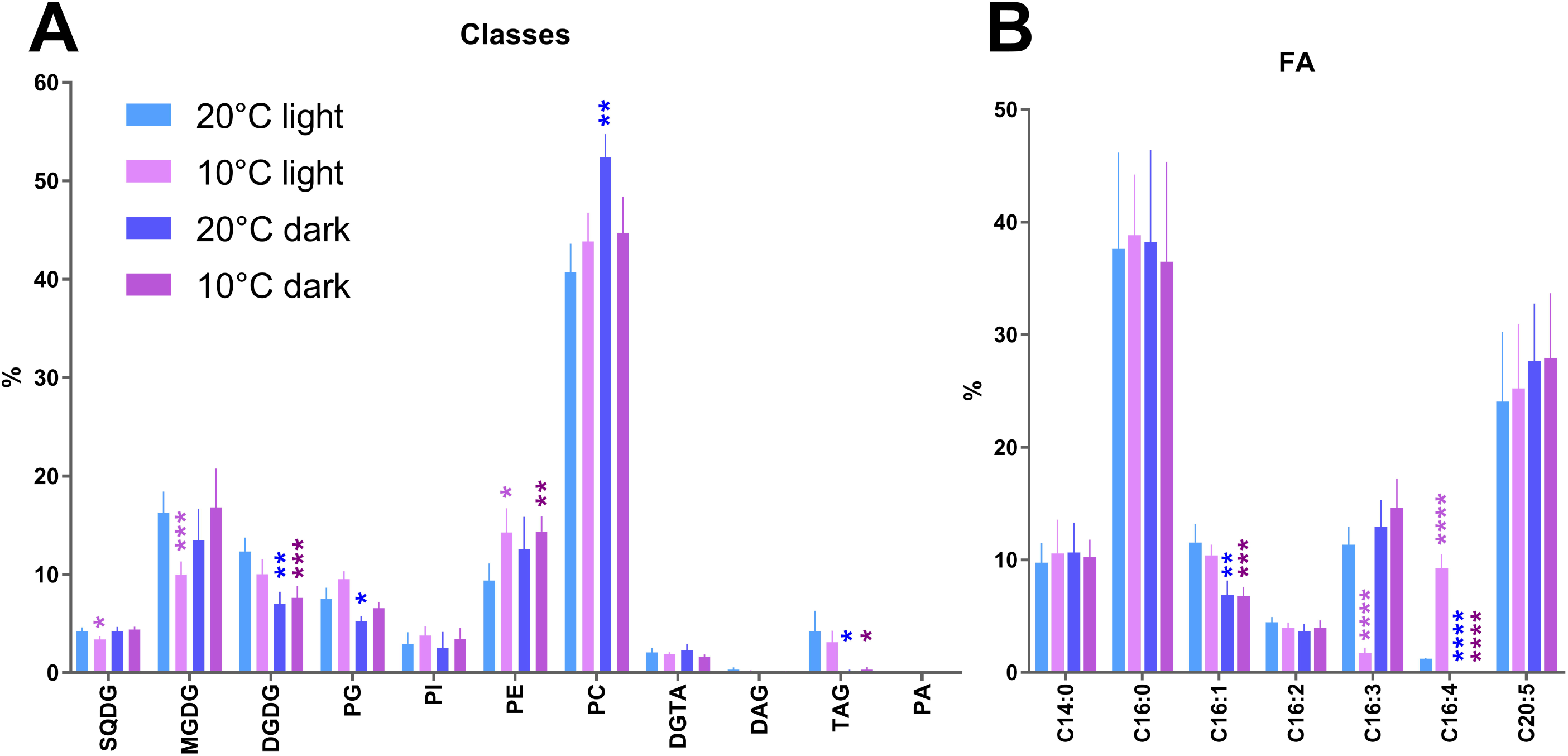
Impact of exposure to 10 °C, 4 days in the darkness (0 µmol photons·m^-2^·s^-1^) and a combination of both on relative proportions (nmol %) of lipid classes (A) and fatty acids (B) compared to 20 °C light:dark (L:D) photoperiod and 40 µmol photons·m^-2^·s^-1^ irradiance. Histograms represent the mean and the error bars the standard deviation of 6 biological replicates. Statistical significance determined using the Bonferroni-Dunn method, with alpha = 0.05. Adjusted p-values: * = p<5×10^−2^, ** = p<1×10^−3^, *** = p<1×10^−4^, **** = p<1×10^-5^.

Total fatty acid profile showed mild variations across the three experimental conditions tested (Fig. 1B). The most notable and statistically significant changes were observed in the cold-exposed cultures in 12:12 L:D photoperiod, characterized by a decrease in 16:3 and a concomitant increase in 16:4. The increase of the proportion of 16:4 critical for plastid glycerolipid lateral fluidity (Dolch and Maréchal, 2015), thus requires an active photosynthesis process or a light-activated pathway. In both dark conditions 16:3 slightly increased although non significantly (Fig. 1B) and 16:4 totally disappeared, in accordance with the downregulation of the putative desaturase producing it (Matthijs *et al*., 2017). Additionally, a reduction in 16:1 was observed under both dark conditions (Fig. 1B).

The level of fatty acid unsaturation was quantified using the double bond index (DBI), which reflects the average number of unsaturations across all molecular species. Noteworthy, the total fatty acid DBI did not vary among the different conditions (Supplementary Figure S1), nevertheless calculating the DBI within each lipid class, some variations were recorded (Fig. 2). SQDG emerged as the least unsaturated plastidial lipid (Fig. 2). Its DBI increased by 17 % across the three experimental conditions, varying from 1.2 to 1.4. A similar percentage increase was observed for MGDG at 10 °C, where the DBI rose from 6.4 to 7.5, whereas a smaller increase of approximately 6 % was noted under both dark conditions. The unsaturation level of DGDG increased by 6 % under low-temperature conditions and by 17 % under both dark conditions. In contrast, PG was the only plastidial lipid to show a reduction in DBI (15 %) at low temperature, but its DBI increased by 25 % and 20 % under dark and dark-cold conditions, respectively. The responses of extraplastidial phospholipids, however, were inconsistent across the experimental treatments.

**Figure 2.**
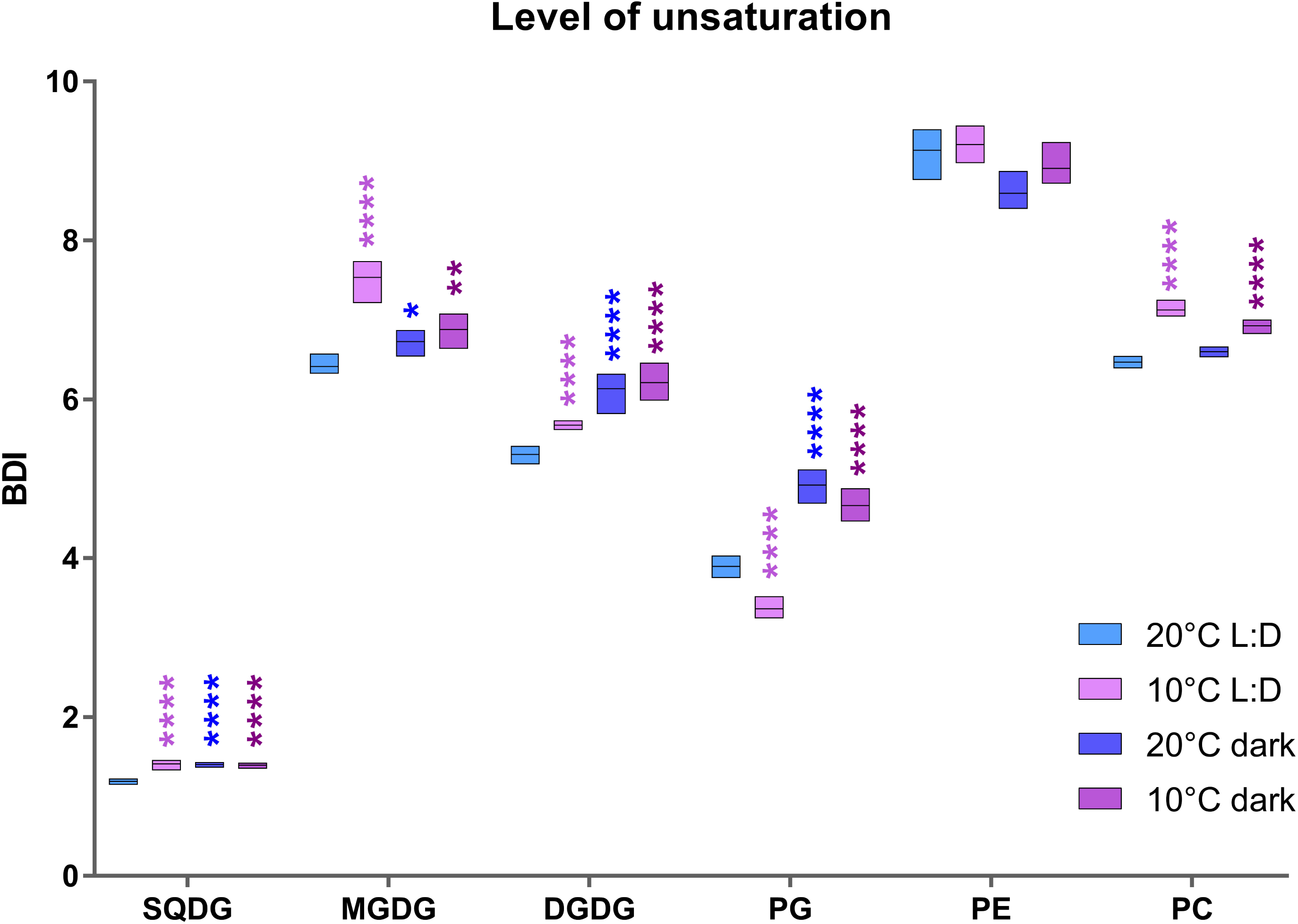
Impact of exposure to 10 °C, 4 days in the darkness (0 µmol photons·m^-2^·s^-1^) and a combination of both on level of unsaturation measured by double bond index (DBI) of major lipid classes compared to 20 °C light:dark (L:D) photoperiod and 40 µmol photons·m^-2^·s^-1^ irradiance. Box plots represent 6 (5 for the 20 °C dark condition) biological replicates. Statistical significance determined using the Bonferroni-Dunn method, with alpha = 0.05. Adjusted p-values: * = p<5×10^−2^, ** = p<1×10^−3^, *** = p<1×10^−4^, **** = p<1×10^-5^.

### Distinct effects of temperature decrease on plastidial lipid species, under control photoperiod conditions and long-exposure to darkness

Under 12:12 L:D photoperiod conditions, galactoglycerolipids MGDG and DGDG showed a global increase of highly unsaturated species (Figs 3A-B) with MGDG-32:7 (mainly composed of MGDG-16:4-16:3) and MGDG-36:9 (mainly composed of MGDG-20:5-16:4) increasing by more than five- and sevenfold, respectively (Fig. 3A). This striking increase in 32:7 was at the expense of 32:4 and 32:5 molecular species, used as substrates for desaturation. Likewise, the increase in 36:9 was at the expense of 36:7 and 36:8. Despite its weak unsaturation, the relative abundance of MGDG-32:1 remained unchanged when cells were cultured at 10 °C.

**Figure 3.**
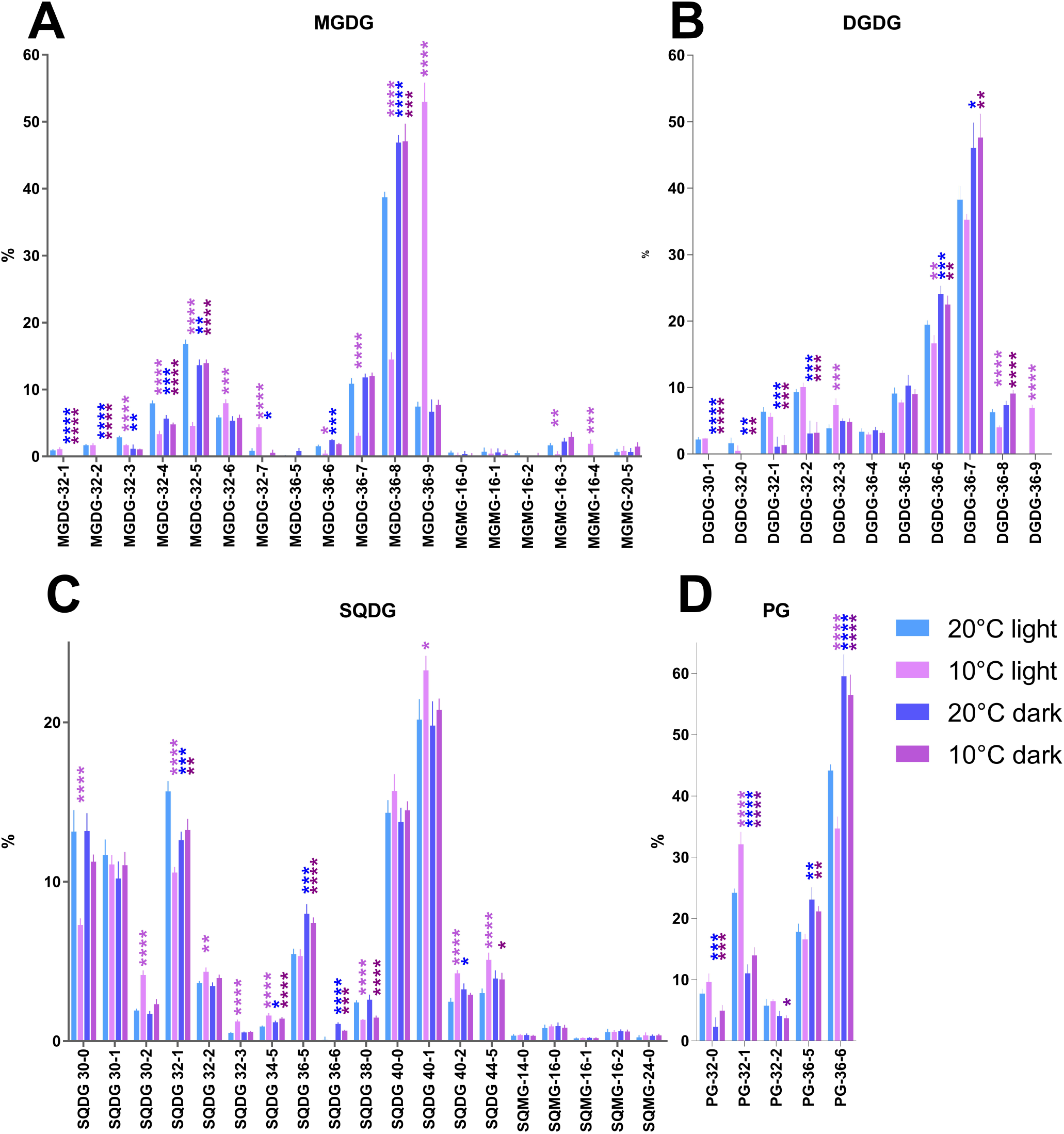
Impact of exposure to 10 °C, 4 days in the darkness (0 µmol photons·m^-2^·s^-1^) and a combination of both compared to 20 °C light:dark (L:D) photoperiod and 40 µmol photons·m^-2^·s^-1^ irradiance on relative proportions (nmol %) of A) MGDG, B) DGDG, C) SQDG, D) PG lipid molecular. A vertical dotted line separates MGDG from MGMG (A) and SQDG from SQMG (C). Histograms represent the mean and the error bars the standard deviation of 6 (5 for the 20 °C dark condition) biological replicates. Statistical significance determined using the Bonferroni-Dunn method, with alpha = 0.05. Adjusted p-values: * = p<5×10^−2^, ** = p<1×10^−3^, *** = p<1×10^−4^, **** = p<1×10^-5^.

Under extended periods of darkness, no increase in 32:7 or 36:9 was observed, nor were there detectable changes in their respective substrates, 32:4 and 32:5 for the former, and 36:7 and 36:8 for the latter. This confirms the requirement of light for the bulk increase in 16:4 species. Low-temperature stress is known to induce changes in the lipidome of plants, including an increase in monogalactosylmonoacylglycerol (MGMG) levels, which result from the hydrolytic cleavage of the ester bond linking an acyl chain to the glycerol backbone of monogalactosyldiacylglycerol (MGDG). In the centric diatom *Stephanodiscus* sp. fluctuations in the levels of MGMG-16:3 were observed when cells were exposed to low temperatures (referred to as lyso-MGDG in Chen *et al*. (2013)), hence we analyzed the main MGMG species. Indeed, MGMG-16:3 and MGMG-16:4 varied in opposite direction (Fig. 3A), as expected from the FA profiles (Fig. 1B) and the decrease of MGDG-36:8 (exclusively composed of MGDG-20:5-16:3) and the increase of MGDG-36:9 (exclusively composed of MGDG-20:5-16:4) (Fig. 3A). This dynamic production of MGMG-16:4 was coherently observed under 12:12 L:D photoperiod, but not in long darkness condition.

Conversely, for DGDG species (Fig. 3B), a slight increase of DGDG-32:3 (mainly represented by DGDG-16:1-16:2) and a mild decrease of DGDG-36:6 (composed of DGDG-20:5-16:1) were recorded, accompanied by the appearance of DGDG-36:9 (most probably composed of DGDG-20:5-16:4), under 12:12 L:D photoperiod conditions. It is highly likely that the appearance of DGDG-36:9 is due to the striking sevenfold increase of the substrate of the DGDG-synthase involved in its production. No such remodeling in DGDG molecular species was observed in dark condition.

SQDG, one of the most species-rich lipid in the plastid (Abida *et al*., 2015), showed an increase of all the species containing 16:2, 24:0 and 20:5 (Fig. 3C). PtFAD6 (Phatr3_J48423, annotated as PTD12) is the possible enzyme responsible for the desaturation of 16:1 to 16:2 (Domergue *et al*., 2003) and it was hypothesized to act on MGDG, DGDG as well as on SQDG (Dolch and Maréchal, 2015). Our results concur with this hypothesis, at least at 10 °C. The observed increase in 20:5-containing SQDG aligns with the low-temperature-induced enhancement of unsaturation. However, the mechanism leading to the increase in 24:0-containing SQDG species remains unclear. 24:0 is an enigmatic fatty acid, for which at the time of writing the synthetic pathway and function in *P. tricornutum* is not known. Two Sulfoquinovosylmonoacylglycerol (SQMG) species were reported as biomarkers in the centric diatom *Stephanodiscus* sp. under cold stress (referred to as lyso-SQDG in Chen *et al*. (2013)), but in our experiments, none of the SQMG species showed a statistically significant variation among the conditions tested (Fig. 3C).

Phosphatidylglycerol (PG) is the sole phospholipid synthesized within plastids and highly likely localizing to the thylakoids (Svenning *et al*., 2024). Among PG species, two distinct pools can be identified, plastidial and extraplastidial. A statistically significant decrease in PG-36:6 (composed exclusively of PG-20:5-16:1) was observed, while PG-32:1 (PG-16:1-16:0) increased. It is highly likely that the 16:1 fatty acid esterified at the *sn-*2 position of PG-36:6 contains a *trans* double bond at position Δ3, introduced by PtFAD4 (Phatr3_J5271, referred to as Phatr_41301 in Dolch and Maréchal (2015)). PtFAD4 appears to be insensitive to cold activation. In higher plants, PG molecular species containing saturated and *trans*-unsaturated fatty acids are more prevalent in chilling-sensitive species, leading to high phase transition temperatures and a predominance of gel-phase domains at lower temperatures (Sklar *et al*., 1979). This phenomenon directly correlates with increased membrane rigidity and reduced functionality under cold stress conditions. In diatoms, PG composition adjusts dynamically in response to temperature shifts (Fig. 3D). Given that the introduction of a *cis*-unsaturated bond in a PG molecule substantially lowers its phase transition temperature, an increase in PG species containing 16:1 with a *cis* double bond at low temperatures would enhance membrane fluidity. Conversely, PG species containing 16:1 with a *trans* double bond would decrease, as *trans*-unsaturated fatty acids behave more like saturated fatty acids because of its stiffness, favoring gel-phase formation (Murata and Yamaya, 1984).

#### Light is required for plastidial membrane lipid homeoviscous adaptation to low temperature

The response of plastidial lipids (MGDG, DGDG, SQDG, and PG) to low temperature upon long exposure to darkness appears as moderate, with a few notable exceptions, such as SQDG-38:0 (exclusively composed of SQDG-14:0-24:0), which in the dark at 10 °C aligns more closely with the trend observed in the 12:12 L:D photoperiod condition at 10 °C (Fig. 3C). The primary distinctions between the two dark treatments and the 12:12 L:D photoperiod condition at 10 °C are observed in MGDG-36:8 (Fig. 3A), as well as PG species PG-32:0, PG-32:1, PG-36:5, and PG-36:6 (Fig. 3D), which show opposing trends. These findings suggest that light is required for plastidial lipid acclimation to low temperature. One could speculate that plastids and their associated metabolic processes are more profoundly affected by the absence of light than by low-temperature exposure. It is also possible that the homeoviscous adaptation of plastid membrane is needed only upon light exposure, and therefore controlled by light, whereas this costly acclimation may not be triggered in long exposure to darkness (mimicking descent or sinking in the water column).

### Cold stress elevates PC and PE unsaturation under photoperiodic and extended dark conditions

PC and PE are the predominant phospholipids in the endomembrane system of *P. tricornutum*, with PC displaying the greatest species diversity; in contrast, PE species are characterized by the presence of 20:5 at the *sn-*1 position (Abida *et al*., 2015). A notable decrease in PC species containing C16 and C18 FA with fewer than three unsaturations was observed under reduced temperatures, such as the nearly 50% reduction of PC-32:0 at 10 °C compared to 20 °C (Fig. 4A). Conversely, species containing VLC-PUFAs increased significantly, including a more than twofold rise in PC-42:11 (Fig. 4B). PE species were affected differently compared to PC (Fig. 4C). Among the 36C containing species, only PE-36:6 (composed of PE-20:5-16:1) presented an almost 28 % reduction (from 10 % in 20 °C light condition to 7.23 % at 10 °C). Surprisingly, PE-40:9 and PE-40:10 showed a reduction although being composed of VLC-PUFAs at both *sn-*1 and *sn-*2 positions. On the contrary, PE-42:11 (PE-20:5-22:6) showed a relative increase of 28 %, statistically significantly rising from 25 % at 20 °C to 32 % at 10 °C (Fig. 4C). The observed increase might possibly be due to an increased activity of PtELO5a (Phatr3_J9255) induced from the drop of temperature.

**Figure 4.**
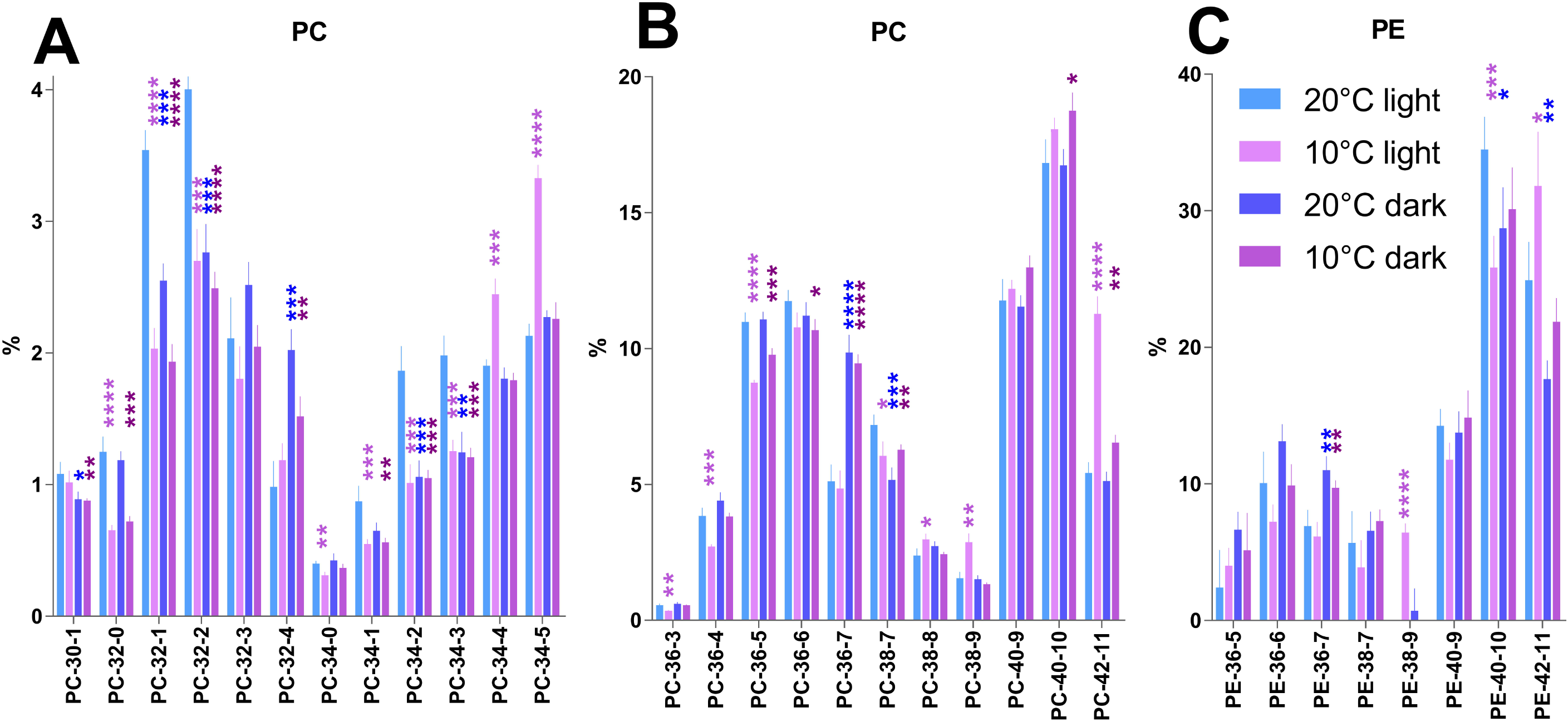
Impact of exposure to 10 °C, 4 days in the darkness (0 µmol photons·m^-2^·s^-1^) and a combination of both compared to 20 °C light:dark (L:D) photoperiod and 40 µmol photons·m^-2^·s^-1^ irradiance on relative proportions (nmol %) of A) PC (low abundance species), B) PC high abundance species), C) PE lipid molecular species. Histograms represent the mean and the error bars the standard deviation of 6 (5 for the 20 °C dark condition) biological replicates. Statistical significance determined using the Bonferroni-Dunn method, with alpha = 0.05. Adjusted p-values: * = p<5×10^−2^, ** = p<1×10^−3^, *** = p<1×10^−4^, **** = p<1×10^-5^.

PtELO5a is a Δ5 elongase involved in the synthesis of 22:6 (Zhu *et al*., 2024). Knockout mutants of this gene showed reduced PE-42:11 and increased levels of PE-40:9 and PE-40:10 along with a hypersensitivity to high temperatures (28 °C, Zhu *et al*., 2024). This finding could provide new insights into the mechanisms cells employ to adapt to both high and low temperature conditions. Along with the variations described above, PE-38:9 (exclusively composed of PE-20:5-18:4) showed an impressive increase from undetectable levels at 20 °C to almost 6.5 % at 10 °C (Fig. 4C). This finding is in line with the increase of the unsaturation level in membrane lipids to cope with low temperatures.

### Endomembrane lipid homeoviscous adaptation to low temperatures is less dependent on light

PC and PE species overall responded differently to the two dark conditions and in general, the dark condition at 10 °C followed the 12:12 L:D photoperiod condition at 10 °C (Fig. 4). A few exceptions are noteworthy, PC-34:2 PC-34:3, decreased in all the tested conditions, whereas PC-34:4 and PC-34:5 did not show any variation in the dark compared to the control whereas at 10 °C in the light both statistically significantly increased (Fig. 4A). The most striking exception is PC-36:7 (exclusively composed of PC-20:5-16:2) that almost doubled in both dark conditions (Fig. 4B).

## Discussion

The diatom *Phaeodactylum tricornutum*, has been employed here as a model organism to investigate lipidomic responses to decreasing temperatures and extended periods of darkness, such as those encountered when descending in the water column in Atlantic or Pacific oceanic contexts (but see Hoppe *et al*. (2024) for a more accurate description of the extent of the euphotic zone). Reports on the lipidomic response of *P. tricornutum* to low temperatures remain scarce (Thompson *et al*., 1992*a*,*b*; Jiang and Gao, 2004; Fierli *et al*., 2022; Dorrell *et al*., 2024), particularly those examining lipid classes and molecular species in detail. In contrast, more comprehensive studies have been conducted on certain eustigmatophytes, unicellular algae within the same monophyletic SAR supergroup (Gill *et al*., 2018; Willette *et al*., 2018; Carneiro *et al*., 2020; Chua *et al*., 2020). In plants, the study of freezing (sub-zero temperatures) and chilling (above-zero temperatures) tolerance is driven by the need to cultivate economically significant crops across diverse latitudes (Moellering *et al*., 2010; Grossman, 2023). In diatoms, however, the focus on low-temperature resistance is rooted in understanding their ecological success and remarkable resilience. This divergence in research priorities supports the substantial disparity in knowledge between plants and diatoms regarding adaptation to cold environments.

In plants, cold adaptation is associated with an increase in phospholipid content and the unsaturation level of fatty acids (Steponkus, 1984), whereas freezing tolerance induces a remodeling of chloroplast membrane lipids (Li *et al*., 2008) including the synthesis of polygalactosyldiacylglycerols (triGDG, tetraGDG) through the action of the product of the SENSITIVE TO FREEZING2 (SFR2) locus which encodes for a galactolipid:galactolipid glycosyltransferase (GGGT). SFR2 gene product catalyzes the transfer of a galactose moiety from a MGDG molecule to another in a processive fashion (van Besouw and Wintermans, 1978; Benning and Ohta, 2005). The role of SFR2-related processes, including the production of tri- and tetraGDG is currently comprehended in contexts where cells may be disrupted by intracellular ice crystal formation.

Although freezing conditions is not addressed here, SFR2 homologs were searched in the *P. tricornutum* genome, but no hits were found, in agreement with the absence of triGDG or tetraGDG in the published glycerolipidomes.

The main response to low temperature was an increase in phospholipid and concomitant decrease in MGDG and DGDG (Fig. 1A) along with an almost 85% reduction of 16:3 counterbalanced by a 5.2-fold increase of 16:4. A reanalysis of lipidomics data reported in Supplementary Dataset S6 produced by Dorrell *et al*. (2024), corroborated most of our findings, although substantial differences in the cultivation conditions are present. Namely, Dorrell and colleagues have exposed *P. tricornutum* cultures to 8 °C without agitation and in a continuous light regime. The control conditions were either 19 °C in 12:12 L:D photoperiod or 19 °C in continuous light. In Supplementary Figures S2-S4, we report the results of such reanalysis. The galactolipid reduction was not observed at all by Dorrell *et al*. (2024) in low temperature (Supplementary Fig. S2A), conversely, the unbalance between 16:3 and 16:4 (Supplementary Fig. S2B) (possibly erroneously identified as 18:0) was validated. In GC-FID analyses, the retention times of 18:0 and 16:4 are indistinguishable, making it impossible to differentiate between them. However, mass spectrometry revealed that the lipidome of *P. tricornutum* contains only traces of 18:0 (present paper, Abida *et al*., 2015). Consequently, we deduced that the ’18:0 or 16:4’ peak observed in GC-FID from cold-exposed cultures predominantly represents 16:4. This interpretation is consistent with the reanalyzed dataset reported by Dorrell *et al*. (2024) (Supplementary Fig. S2B). The mentioned 16:3-to-16:4 switch in response to dropping temperatures was also recorded (Thompson *et al*., 1992*b*).

The MGDG:DGDG ratio is a key determinant of the structural and functional integrity of chloroplast membranes (Chng *et al*., 2022). These two galactolipids constitute the bulk of thylakoid membranes and play distinct yet complementary roles in membrane dynamics. MGDG, a non-bilayer-forming lipid due to its cone-shaped molecular geometry, contributes to membrane curvature and flexibility, whereas DGDG, a bilayer-forming lipid, stabilizes membrane structure by promoting lamellar phases (Garab *et al*., 2016). An optimal MGDG:DGDG ratio is essential to maintain the delicate balance between membrane fluidity and stability, particularly under stress conditions or during development (Shimojima and Ohta, 2022).

Importantly, this lipid ratio is not static but undergoes dynamic remodeling in response to environmental changes, especially temperature fluctuations. When temperatures drop, plants and algae often respond by decreasing the MGDG:DGDG ratio to preserve membrane lamellarity and prevent phase separation or fusion events that could destabilize the thylakoid structure. In this context, an excess of MGDG, which favors the formation of hexagonal II phases, can compromise chloroplast compartmentalization by inducing non-lamellar transitions and promoting membrane fusion.

Our results show that the MGDG:DGDG ratio favored DGDG at 10 °C and in 12:12 L:D photoperiod (Fig. 5). A shift toward higher DGDG content under cold stress reinforces the bilayer structure, enhancing membrane stability and maintaining photosynthetic efficiency. These findings highlight that the MGDG:DGDG ratio functions as a critical biophysical and regulatory parameter of membrane homeostasis. Its temperature-dependent tuning is part of a broader adaptive strategy to preserve membrane integrity, maintain thylakoid architecture, and sustain photosynthetic activity under adverse environmental conditions. This effect is not observed in the dark as there is no photosynthetic activity.

**Figure 5.**
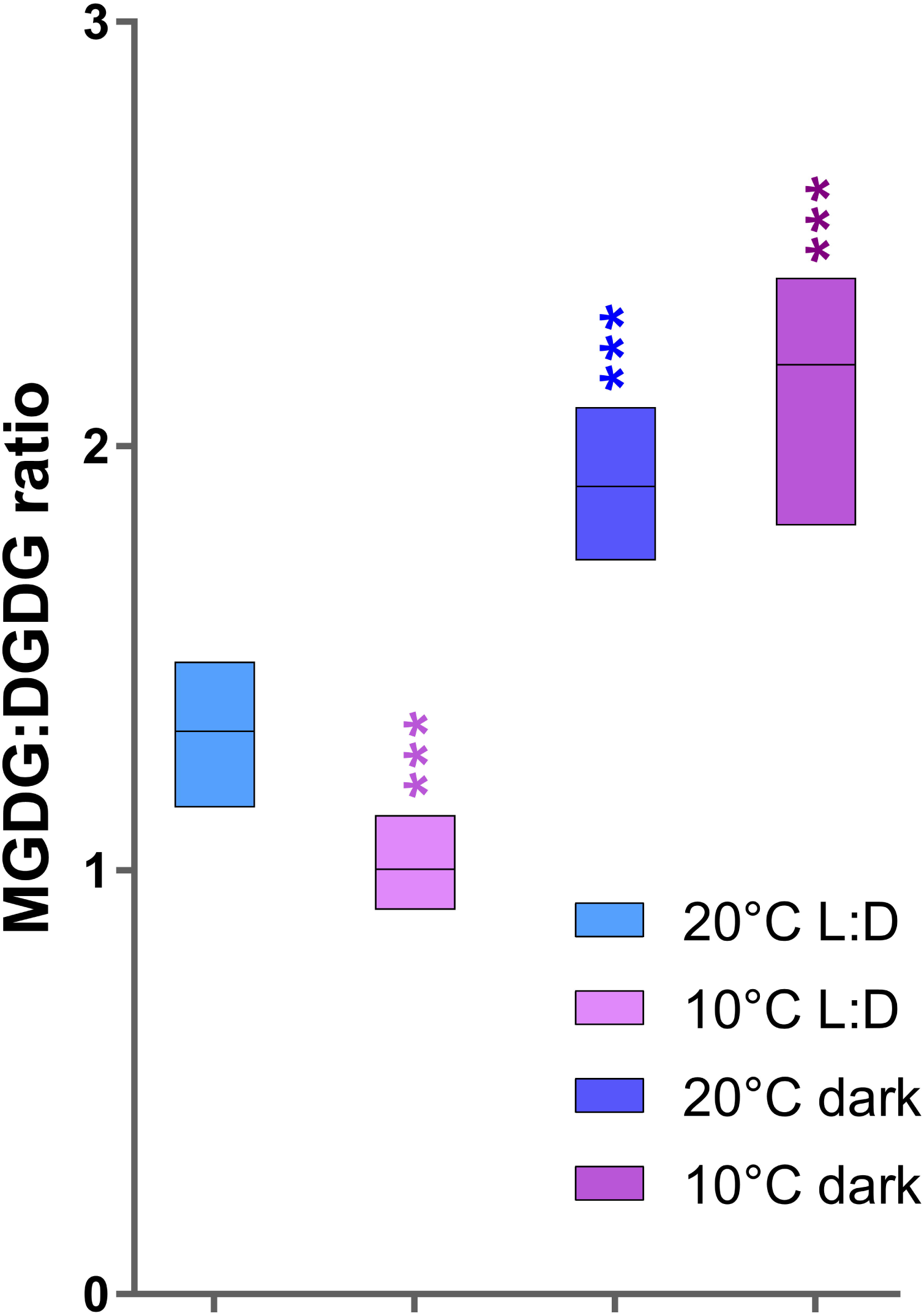
Impact of exposure to 10 °C, 4 days in the darkness (0 µmol photons·m^-2^·s^-1^) and a combination of both compared to 20 °C light:dark (L:D) photoperiod and 40 µmol photons·m^-2^·s^-1^ irradiance on MGDG:DGDG ratio. Box plots represent 6 (5 for the 20 °C dark condition) biological replicates. Statistical significance determined using the Bonferroni-Dunn method, with alpha = 0.05. Adjusted p-values: * = p<5×10^−2^, ** = p<1×10^−3^, *** = p<1×10^−4^, **** = p<1×10^-5^.

Overall, it has been demonstrated that lipid unsaturation levels in oceanic phytoplankton are correlated to the water temperature (Holm *et al*., 2022) and such a general trend is validated in our results (Fig. 2), although no visible increase in 20:5 was recorded (Fig. 1B). A 2-fold increment in 20:5 percentage was recorded in *P. tricornutum* cells shifted from 25 °C to 10 °C (Jiang and Gao, 2004) and a non-significant increase when shifted from 20 °C to 10 °C (Fierli *et al*., 2022), possibly revealing that the 20-to-10 °C shift we have imposed to our cultures was not stressing enough to induce 20:5 variations. Nevertheless, Dorrell *et al*. (2024) recorded a statistically significant 13 % decrease of 20:5 in cells exposed to 8 °C (continuous light) when compared to cells exposed to 19 °C 12:12 L:D photoperiod and an almost 22 % increase when compared to cells exposed to 19 °C continuous light (Supplementary Fig. S2B). More generally, the PUFA percentage in different phytoplankton species show a trend to increase with lowering temperatures, but this is not a general rule (Mortensen *et al*., 1988; Thompson *et al*., 1992*b*). Notably, as already demonstrated in other systems, desaturases are activated by low temperatures (Tiku *et al*., 1996; Domínguez *et al*., 2010; An *et al*., 2013; Ishikawa *et al*., 2024) and *P. tricornutum* desaturases are no exception (Fierli *et al*., 2022). In the eustigmatophytes *Nannochloropsis oceanica* (Carneiro *et al*., 2020; Chua *et al*., 2020) and *Microchloropsis salina* (referred to as *Nannochloropsis salina* in Gill *et al*. (2018), Willette *et al*. (2018)) the reported trends for 20:5 across temperature ranges are inconsistent, with levels either increasing, decreasing, or remaining stable. It is noteworthy that culture conditions differed substantially compared to ours because in three out of the four studies, photobioreactors (larger culture volume) and much higher irradiances were used. This variability shows the complex and species-specific nature of fatty acid regulation, reflecting potentially diverse adaptive strategies to environmental temperature fluctuations. Again, none of the studies reported above, measured 16:3 and 16:4, although eustigmatophytes contain much lower proportions of these FAs (Billey *et al*., 2021).

To the best of our knowledge, no targeted lipidomics approach has yet been applied to study the effects of low-temperature and darkness on *P. tricornutum* (but see Dorrell *et al*. (2024) and the reanalysis performed here, supplementary Figs S2-S4). Such an approach, however, has the potential to reveal molecular lipid species most affected by these conditions.

One of the most striking effects of temperature decrease was observed in MGDG-36:9, which exhibited a more than seven-fold increase at 10 °C compared to 20 °C, accompanied by a significant reduction in the predominant MGDG species, MGDG-36:8 (Fig. 3A, Supplementary Fig. S3A). From the lipidomics analyses of a *P. tricornutum MGD*_γ_ KO mutants (Guéguen *et al*., 2024), it can be hypothesized that MGDγ produces MGDG-36:9, possibly revealing a role played by this enzyme in the response to low temperatures to tune the highly unsaturated MGDG species. A similar variation occurred in MGDG-32:7 and MGDG-32:6 which became the two dominant species, accounting altogether for 50 % of all the MGDG-32:x species at 10 °C, compared to 20 °C where they accounted for only 16% of total double C16 MGDG species. This shift in the dominant MGDG species in low-temperature conditions reflects an increase in 16:4 at the expense of 16:3. This trend is further supported by similar changes in MGMG species, with MGMG-16:3 decreasing and MGMG-16:4 increasing. The polyunsaturated to less saturated fatty acid ratio is tightly correlated to membrane fluidity in low-temperature conditions and in different photosynthetic organisms it stands true for chloroplast membranes as well (Routaboul *et al*., 2000; Morgan-Kiss *et al*., 2006). To maintain efficient and properly folded photosystems, high levels of unsaturation in membrane lipids are crucial (Seiwert *et al*., 2017; Zorin *et al*., 2017). Temperature downshifts affect membrane fluidity. The distinct responses of plastidial lipids under the two dark conditions compared to low temperature condition in light suggest that because the absence of light does not influence photosystems, the lipid regulation systems remain unresponsive. This implies a finely tuned regulation of plastidial lipids, potentially adjusting the MGDG:DGDG ratio and species composition to adapt to external environmental conditions.

DGDG species did not vary much at 10 °C in photoperiodic conditions compared to 20 °C (with the exception of the appearance of DGDG-36:9, Fig. 3B, Supplementary Fig. S3B). The absence of variation in the other species corroborates the hypothesis that most of the plastidial desaturases in *P. tricornutum* primarily target MGDG (Dolch and Maréchal, 2015; Guéguen *et al*., 2024). Nevertheless, it was also hypothesized that PtFAD6 might desaturate 16:1 esterified at DGDG *sn-*1 and *sn-*2 positions. The results presented above, however, only partially support such hypothesis, at least at 10 °C. Specifically, the increase in DGDG-32:3 is not accompanied by a corresponding decrease in its precursor, DGDG-32:2. Similarly, the reduction in DGDG-36:6 does not go with an increase in DGDG-36:7.

In both dark conditions, plastidial lipids show an increase in the level of unsaturation measured through the double bond index (DBI, Fig. 2). This may indicate a somehow accelerated plastidial desaturation with the PtFAD6 up-regulated by +3 log_2_FC in the dark (Matthijs *et al*., 2017; Villar *et al*., 2025). In the dark, photosynthesis reduces but the energy conversion stays active during even long dark periods (Nymark *et al*., 2013; Lacour *et al*., 2019; Joli *et al*., 2024), which can induce an increase in oxidative stress that in turn can be balanced by the increase of unsaturated molecular species. Although *PtPAD* is up-regulated in darkness (+2 log_2_FC Matthijs *et al*., 2017; Villar *et al*., 2025), levels of 16:1 decreased by 40% under both dark conditions (Fig. 1B, Supplementary Fig. S2B). This observation, combined with the stability of 20:5 and the predominance of 16:0 over 16:1 in *P. tricornutum* under standard conditions, suggests that 16:0 is rapidly incorporated into galactolipids and exported faster than it is processed by PtPAD. The differential behavior of 16:1 and 20:5 further implies that 20:5 biosynthesis is more complex than previously assumed and may involve distinct pathways relying on different precursors.

Therefore, it can be hypothesized that the increase in the DBI recorded in the dark at both 10 °C and 20 °C (Fig. 2) is attributable to a prolonged retention of fatty acids within plastidial lipids, where desaturases are available to act on their substrates.

Diatoms inhabiting polar regions face the dual challenges of prolonged darkness and extreme cold, necessitating energy-efficient adaptations for survival. In the polar diatom *Fragilariopsis cylindrus*, autophagy is triggered after three days in the dark (Joli *et al*., 2024), enabling the degradation of lipid droplets via lipophagy, as demonstrated in *P. tricornutum* using KO mutants (Leyland *et al*., 2024). This process may explain the very low levels of TAGs in dark conditions and the reduction in 16:1 (Fig. 1A, B). Lipophagy is critical for recycling lipid droplets into substrates for β-oxidation, sustaining basal metabolic demands during periods of halted photosynthesis.

Diatoms, like any other phytoplankter, live in three-dimensional world characterized by great variability of physico-chemical features. Salinity, density, temperature gradients are highly influenced by turbulence which is produced by a series of sources like, *e.g.* wind, currents, rain, heating. The upper layer of the ocean is characterized by intense mixing due to the abovementioned turbulence and this has led Sverdrup to postulate his critical depth hypothesis (Sverdrup, 1953) that posits that phytoplankton can accumulate when the depth of the mixed layer allows sufficient residence time in the euphotic zone for cells to acquire enough solar energy, enabling growth by cell division to exceed losses due to mixing. Since its publication in 1953, the Sverdrup’s hypothesis has been reviewed thousands of times and more complex hypotheses arose based on the Sverdrup’s critical depth hypothesis. Nevertheless, the concept of phytoplankton cells lost through mixing or turbulence stayed intact (*e.g.* Franks, 2015).

Although richer in nutrients, deeper ocean layers are darker and colder, requiring virtually every diatom cell to endure cold and dark conditions at some point. The results presented here demonstrate that *P. tricornutum* and potentially diatoms in general, can survive to variably long dark periods and low temperatures by activating cellular responses *e.g.* adjustments in lipid and fatty acid composition, morphological and structural modifications, and variations in pigment content (present work; Thompson *et al*., 1992*a*,*b*; Jiang and Gao, 2004; Fierli *et al*., 2022; Dorrell *et al*., 2024; Joli *et al*., 2024). The results presented here suggest that *P. tricornutum* activates a cold response involving the 16:3-to-16:4 fatty acid switch only under illuminated conditions (Fig. 6). While this response may represent a species-specific adaptation of *P. tricornutum*, it is worth considering that other diatom species, particularly those adapted to polar environments, might employ distinct mechanisms to cope with low temperatures. Notably, the absence of light appears to decouple the homeoviscous membrane response. However, this conclusion should be approached with caution, as the minimum irradiance required to sustain photosynthesis is remarkably low (Raven *et al*., 2000). Indeed, it has been demonstrated that polar diatoms can perform photosynthesis even under extremely low irradiance levels (Hoppe *et al*., 2024, and literature therein). This implies that, even below the euphotic zone, the response to temperature reduction observed under the 10 °C 12:12 light:dark cycle may still be relevant.

**Figure 6.**
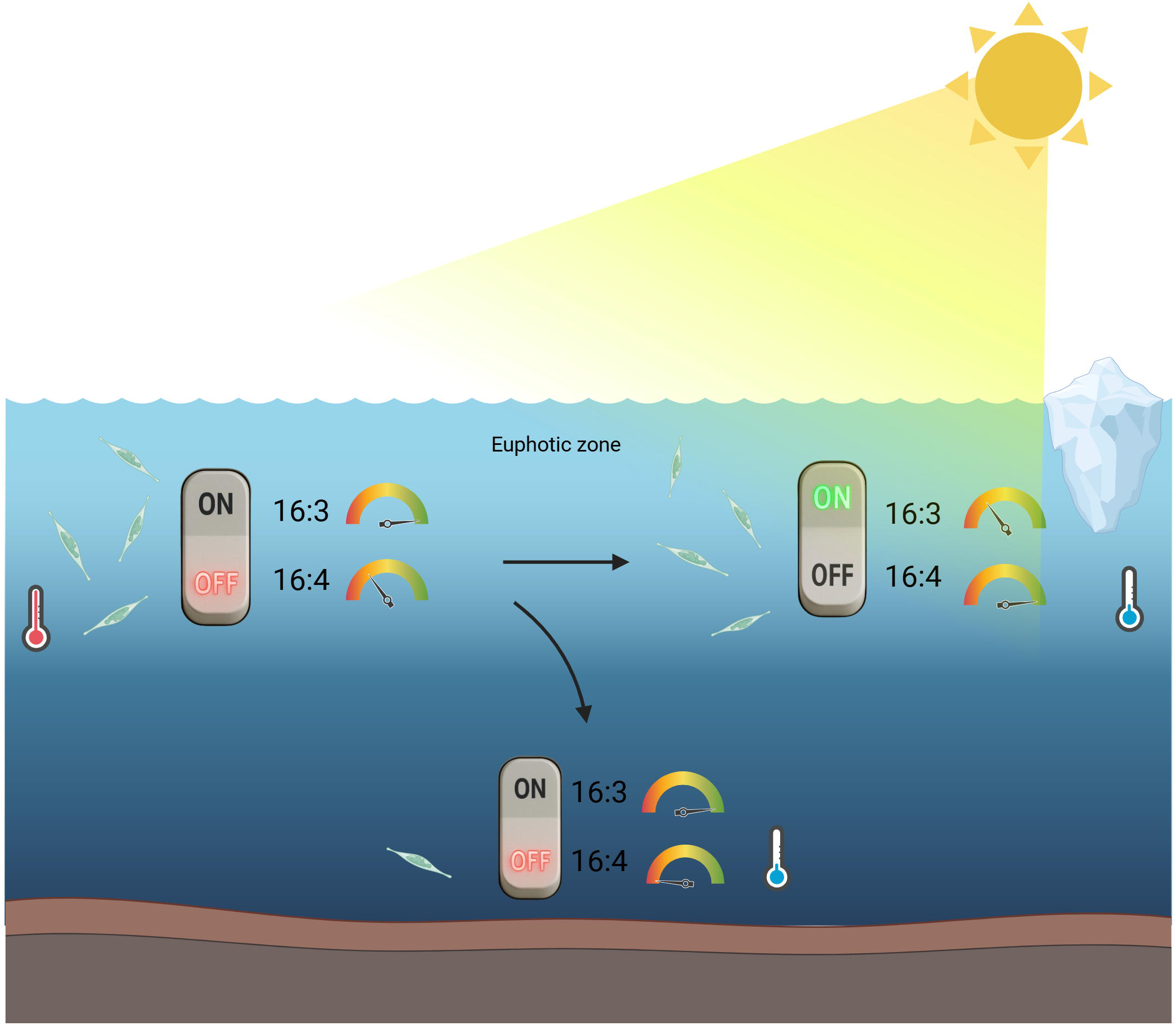
Schematic representation of the decoupling of the homeoviscous response to temperature decrease in the presence or absence of light. Under optimal growth conditions (20 °C, 12:12 L:D), 16:3 is abundant, while 16:4 is nearly absent. When temperature decreases with light present, a shift occurs, increasing 16:4 at the expense of 16:3 (16:3-to-16:4 switch). In contrast, when both temperature decreases and light is absent, this switch is inactivated, and the relative proportions of 16:3 and 16:4 remain unchanged. Created with BioRender (https://www.biorender.com).

These observations witness the complexity of diatom physiological responses to environmental changes. It has already been demonstrated that diatoms respond very rapidly to stress relief (*e.g.* Domingues *et al*., 2012; Helliwell *et al*., 2021; Zhang *et al*., 2024).

Altogether, this highlights that the ecological success of diatoms may lay in their remarkable resilience, which relies on mechanisms that are still not fully understood (*e.g.* Li *et al*., 2023; Flori *et al*., 2024).

## Conclusions

Our study elucidates the lipidomic responses of the model diatom *P. tricornutum* to low temperatures and extended darkness, revealing significant shifts in plastidial lipid species and fatty acid unsaturation levels. These results show the importance of lipid remodeling, particularly the increased prevalence of unsaturated molecular species, as a crucial mechanism to maintain membrane fluidity and protect photosynthetic systems under stress conditions. The absence of polygalactosyldiacylglycerols, such as triDGD and tetraDGD, and the stability of certain lipid species further demonstrate the distinct cold adaptation strategies employed by diatoms compared to plants.

Our findings also demonstrate that light plays a pivotal role in activating plastidial lipid remodeling, specifically the 16:3-to-16:4 switch, under low-temperature conditions. This light-dependent response, coupled with the limited changes observed in darkness, suggests a finely tuned regulatory mechanism that optimizes resource allocation to acclimation processes based on environmental cues. Moreover, the data indicate that endomembrane lipid remodeling is less dependent on light.

These findings contribute to a broader understanding of how diatoms adapt to fluctuating environmental conditions, such as descending into colder, darker ocean layers. Such adaptations likely contribute to their ecological success and resilience in diverse marine environments. Future studies should explore the molecular pathways driving these lipidomic shifts, including the regulation of desaturase activity and the interplay between plastidial and extraplastidial lipid pools. Furthermore, investigating the lipidome responses of other diatom species, particularly polar taxa, could provide insights into species-specific strategies for coping with extreme environments.

## Acknowledgements

This work was supported by the French National Research Agency (GRAL Labex ANR-10-LABEX-04, EUR CBS ANR-17-EURE-0003, ANR AlpAlga ANR-20-CE02-0020, ANR DIM ANR-21-CE02-0021, PEPR Algadvance A-22-PEBB-0002, Glyco@Alps Cross-Disciplinary Program; Grant ANR-15-IDEX-02). YS was supported by a PhD grant from CEA. CSV was supported by a PhD grant from INRAE (BAP programme). Lipid analyses were performed at the LIPANG platform supported by the Rhône-Alpes Region, the FEDER funds, and French National Research Agency (GRAL Labex ANR-10-LABEX-04, EUR CBS ANR-17-EURE-0003).

## Author contribution

AA conceived the study. AA EM supervised the project. YS CSV RB conducted the experiments. MSc MSi produced lipidomes. JJ supervised the lipidome production. AA wrote the first draft of the manuscript. All the authors contributed to the final version of the manuscript.

---

**Supplementary Figure S1.** Impact of exposure to 10 °C, 4 days in the darkness (0 µmol photons·m^-2^·s^-1^) and a combination of both on the level of unsaturation measured by double bond index (DBI) on total fatty acids compared to 20 °C light:dark (L:D) photoperiod and 40 µmol photons·m^-2^·s^-1^ irradiance. Box plots represent 6 biological replicates. No statistically significant of differences were determined using the Bonferroni-Dunn method.

**Supplementary Figure S2.** Reanalysis of lipidomics data produced by Dorrell *et al*. (2024) in Supplementary dataset S6. A. Lipid classes, B. Fatty acids in *P. tricornutum* WT cultures grown at 19 °C 12:12 L:D photoperiod (LD), 8 °C in continuous light (8C) and 19 °C continuous light (CL). Statistical significance determined using the Bonferroni-Dunn method, with alpha = 0.05. Adjusted p-values: * = p<5×10^−2^, ** = p<1×10^−3^, *** = p<1×10^−4^, **** = p<1×10^-5^.

**Supplementary Figure S3.** Reanalysis of lipidomics data produced by Dorrell *et al*. (2024) in Supplementary dataset S6. A. MGDG, B. DGDG, C. SQDG, D. PG in *P. tricornutum* WT cultures grown at 19 °C 12:12 L:D photoperiod (LD), 8 °C in continuous light (8C) and 19 °C continuous light (CL). Statistical significance determined using the Bonferroni-Dunn method, with alpha = 0.05. Adjusted p-values: * = p<5×10^−2^, ** = p<1×10^−3^, *** = p<1×10^−4^, **** = p<1×10^-5^.

**Supplementary Figure S4.** Reanalysis of lipidomics data produced by Dorrell *et al*. (2024) in Supplementary dataset S6. A. PC, B. PE, C. DGTA in *P. tricornutum* WT cultures grown at 19 °C 12:12 L:D photoperiod (LD), 8 °C in continuous light (8C) and 19 °C continuous light (CL). Statistical significance determined using the Bonferroni-Dunn method, with alpha = 0.05. Adjusted p-values: * = p<5×10^−2^, ** = p<1×10^−3^, *** = p<1×10^−4^, **** = p<1×10^-5^.

## References

Abida H, Dolch L-J, Meï C, et al. 2015. Membrane glycerolipid remodeling triggered by nitrogen and phosphorus starvation in *Phaeodactylum tricornutum*. Plant Physiology 167, 118–136.

Alipanah L, Winge P, Rohloff J, Najafi J, Brembu T, Bones AM. 2018. Molecular adaptations to phosphorus deprivation and comparison with nitrogen deprivation responses in the diatom Phaeodactylum tricornutum. (S Lin, Ed.). PLOS ONE 13, e0193335.

Allen AE, LaRoche J, Maheswari U, Lommer M, Schauer N, Lopez PJ, Finazzi G, Fernie AR, Bowler C. 2008. Whole-cell response of the pennate diatom *Phaeodactylum tricornutum* to iron starvation. Proceedings of the National Academy of Sciences 105, 10438– 10443.

Amari C, Carletti M, Yan S, Michaud M, Salvaing J. 2024. Lipid droplets degradation mechanisms from microalgae to mammals, a comparative overview. Biochimie 227, 19–34.

An M, Mou S, Zhang X, Ye N, Zheng Z, Cao S, Xu D, Fan X, Wang Y, Miao J. 2013. Temperature regulates fatty acid desaturases at a transcriptional level and modulates the fatty acid profile in the Antarctic microalga *Chlamydomonas* sp. ICE-L. Bioresource Technology 134, 151–157.

B-Béres V, Stenger-Kovács C, Buczkó K, Padisák J, Selmeczy GB, Lengyel E, Tapolczai K. 2023. Ecosystem services provided by freshwater and marine diatoms. Hydrobiologia 850, 2707–2733.

Benning C, Ohta H. 2005. Three enzyme systems for galactoglycerolipid biosynthesis are coordinately regulated in plants. Journal of Biological Chemistry 280, 2397–2400.

Benoiston A-S, Ibarbalz FM, Bittner L, Guidi L, Jahn O, Dutkiewicz S, Bowler C. 2017. The evolution of diatoms and their biogeochemical functions. Philosophical Transactions of the Royal Society of London. Series B, Biological Sciences 372, 20160397.

Berges JA, Franklin DJ, Harrison PJ. 2002. Evolution of an artificial seawater medium: Improvements in enriched seawater, artificial water over the last two decades. Journal of Phycology 37, 1138–1145.

van Besouw A, Wintermans JFGM. 1978. Galactolipid formation in chloroplast envelopes. Biochimica et Biophysica Acta (BBA) - Lipids and Lipid Metabolism 529, 44–53.

Billey E, Magneschi L, Leterme S, et al. 2021. Characterization of the Bubblegum acyl-CoA synthetase of *Microchloropsis gaditana*. Plant Physiology 185, 815–835.

Bowler C, Allen AE, Badger JH, et al. 2008. The *Phaeodactylum* genome reveals the evolutionary history of diatom genomes. Nature 456, 239–244.

Brown AP, Slabas AR, Rafferty JB. 2009. Fatty Acid Biosynthesis in Plants — Metabolic Pathways, Structure and Organization. In: Wada H, Murata N, eds. Advances in Photosynthesis and Respiration. Lipids in Photosynthesis. Dordrecht: Springer Netherlands, 11–34.

Busseni G, Rocha Jimenez Vieira F, Amato A, et al. 2019. Meta-omics reveals genetic flexibility of diatom nitrogen transporters in response to environmental changes. Molecular Biology and Evolution 36, 2522–2535.

Carneiro M, Cicchi B, Maia IB, Pereira H, Zittelli GC, Varela J, Malcata FX, Torzillo G. 2020. Effect of temperature on growth, photosynthesis and biochemical composition of *Nannochloropsis oceanica*, grown outdoors in tubular photobioreactors. Algal Research 49, 101923.

Chen D, Yan X, Xu J, Su X, Li L. 2013. Lipidomic profiling and discovery of lipid biomarkers in *Stephanodiscus* sp. under cold stress. Metabolomics 9, 949–959.

Chng C-P, Wang K, Ma W, Hsia KJ, Huang C. 2022. Chloroplast membrane lipid remodeling protects against dehydration by limiting membrane fusion and distortion. Plant Physiology 188, 526–539.

Chua ET, Dal’Molin C, Thomas-Hall S, Netzel ME, Netzel G, Schenk PM. 2020. Cold and dark treatments induce omega-3 fatty acid and carotenoid production in *Nannochloropsis oceanica*. Algal Research 51, 102059.

Conte M, Lupette J, Seddiki K, Meï C, Dolch L-J, Gros V, Barette C, Rébeillé F, Jouhet J, Maréchal E. 2018. Screening for biologically annotated drugs that trigger triacylglycerol accumulation in the diatom *Phaeodactylum*. Plant Physiology 177, 532–552.

Dell’Aquila G, Maier UG. 2020. Specific acclimations to phosphorus limitation in the marine diatom *Phaeodactylum tricornutum*. Biological Chemistry 0.

Dell’Aquila G, Zauner S, Heimerl T, Kahnt J, Samel-Gondesen V, Runge S, Hempel F, Maier UG. 2020. Mobilization and cellular distribution of phosphate in the diatom *Phaeodactylum tricornutum*. Frontiers in Plant Science 11, 579.

Dolch L-J, Maréchal E. 2015. Inventory of fatty acid desaturases in the pennate diatom *Phaeodactylum tricornutum*. Marine Drugs 13, 1317–1339.

Dolch L-J, Rak C, Perin G, et al. 2017. A palmitic acid elongase affects eicosapentaenoic acid and plastidial monogalactosyldiacylglycerol levels in *Nannochloropsis*. Plant Physiology 173, 742–759.

Domergue F, Spiekermann P, Lerchl J, Beckmann C, Kilian O, Kroth PG, Boland W, Zähringer U, Heinz E. 2003. New insight into Phaeodactylum tricornutum fatty acid metabolism. Cloning and functional characterization of plastidial and microsomal Δ12-fatty acid desaturases. Plant Physiology 131, 1648–1660.

Domingues N, Matos AR, Marques da Silva J, Cartaxana P. 2012. Response of the Diatom Phaeodactylum tricornutum to Photooxidative Stress Resulting from High Light Exposure. (S Lin, Ed.). PLoS ONE 7, e38162.

Domínguez T, HernÁndez ML, Pennycooke JC, Jiménez P, Martínez-Rivas JM, Sanz C, Stockinger EJ, SÁnchez-Serrano JJ, Sanmartín M. 2010. Increasing _ω_-3 desaturase expression in tomato results in altered aroma profile and enhanced resistance to cold stress. Plant Physiology 153, 655–665.

Dorrell RG, Zhang Y, Liang Y, et al. 2024. Complementary environmental analysis and functional characterization of lower glycolysis-gluconeogenesis in the diatom plastid. The Plant Cell 36, 3584–3610.

Ernst R, Ejsing CS, Antonny B. 2016. Homeoviscous Adaptation and the Regulation of Membrane Lipids. Journal of Molecular Biology 428, 4776–4791.

Fierli D, Barone ME, Graceffa V, Touzet N. 2022. Cold stress combined with salt or abscisic acid supplementation enhances lipogenesis and carotenogenesis in *Phaeodactylum tricornutum* (Bacillariophyceae). Bioprocess and Biosystems Engineering 45, 1967–1977.

Flori S, Dickenson J, Gaikwad T, Cole I, Smirnoff N, Helliwell KE, Brownlee C, Wheeler GL. 2024. Diatoms exhibit dynamic chloroplast calcium signals in response to high light and oxidative stress. Plant Physiology 197, kiae591.

Flori S, Jouneau P-H, Bailleul B, et al. 2017. Plastid thylakoid architecture optimizes photosynthesis in diatoms. Nature Communications 8, 15885.

Flori S, Jouneau P-H, Finazzi G, Maréchal E, Falconet D. 2016. Ultrastructure of the periplastidial compartment of the diatom *Phaeodactylum tricornutum*. Protist 167, 254–267.

Folch J, Lees M, Sloane Stanley GH. 1957. A simple method for the isolation and purification of total lipides from animal tissues. The Journal of Biological Chemistry 226, 497–509.

Frallicciardi J, Melcr J, Siginou P, Marrink SJ, Poolman B. 2022. Membrane thickness, lipid phase and sterol type are determining factors in the permeability of membranes to small solutes. Nature Communications 13, 1605.

Franks PJS. 2015. Has Sverdrup’s critical depth hypothesis been tested? Mixed layers vs. turbulent layers. ICES Journal of Marine Science 72, 1897–1907.

Gagneux-Moreaux S, Moreau C, Gonzalez J-L, Cosson RP. 2007. Diatom artificial medium (DAM): a new artificial medium for the diatom *Haslea ostrearia* and other marine microalgae. Journal of Applied Phycology 19, 549–556.

Garab G, Ughy B, Goss R. 2016. Role of MGDG and non-bilayer lipid phases in the structure and dynamics of chloroplast thylakoid membranes. Sub-Cellular Biochemistry 86, 127–157.

Garab G, Yaguzhinsky LS, Dlouhý O, Nesterov SV, Špunda V, Gasanoff ES. 2022. Structural and functional roles of non-bilayer lipid phases of chloroplast thylakoid membranes and mitochondrial inner membranes. Progress in Lipid Research 86, 101163.

Geneste T, Faure J-D. 2022. Plant polyunsaturated fatty acids: Biological roles, regulation and biotechnological applications. Advances in Botanical Research. Elsevier, 253–286.

Ghysels A, Krämer A, Venable RM, Teague WE, Lyman E, Gawrisch K, Pastor RW. 2019. Permeability of membranes in the liquid ordered and liquid disordered phases. Nature Communications 10, 5616.

Gill S, Willette S, Dungan B, Jarvis J, Schaub T, VanLeeuwen D, St. Hilaire R, Holguin F. 2018. Suboptimal temperature acclimation affects Kennedy pathway gene expression, lipidome and metabolite profile of *Nannochloropsis salina* during PUFA enriched TAG synthesis. Marine Drugs 16, 425.

Grossman JJ. 2023. Phenological physiology: seasonal patterns of plant stress tolerance in a changing climate. The New Phytologist 237, 1508–1524.

Guéguen N, Sérès Y, Cicéron F, et al. 2024. Monogalactosyldiacylglycerol synthase isoforms play diverse roles inside and outside the diatom plastid. The Plant Cell 36, 5023– 5049.

Harrison PJ, Waters RE, Taylor FJR. 1980. A broad spectrum artificial sea water medium for coastal and open ocean phytoplankton. Journal of Phycology 16, 28–35.

Helliwell KE, Harrison EL, Christie-Oleza JA, et al. 2021. A novel Ca2+ signaling pathway coordinates environmental phosphorus sensing and nitrogen metabolism in marine diatoms. Current Biology 31, 978–989.e4.

Holm HC, Fredricks HF, Bent SM, Lowenstein DP, Ossolinski JE, Becker KW, Johnson WM, Schrage K, Van Mooy BAS. 2022. Global ocean lipidomes show a universal relationship between temperature and lipid unsaturation. Science 376, 1487–1491.

Hoppe CJM, Fuchs N, Notz D, et al. 2024. Photosynthetic light requirement near the theoretical minimum detected in Arctic microalgae. Nature Communications 15, 7385.

Ishikawa T, Domergue F, Amato A, Corellou F. 2024. Characterization of unique eukaryotic sphingolipids with temperature-dependent Δ8-unsaturation from the picoalga *Ostreococcus tauri*. Plant & Cell Physiology 65, 1029–1046.

Jiang H, Gao K. 2004. Effects of lowering temperature during culture on the production of polyunsaturated fatty acids in the marine diatom *Phaeodactylum tricornutum* (Bacillariophyceae). Journal of Phycology 40, 651–654.

Joli N, Concia L, Mocaer K, et al. 2024. Hypometabolism to survive the long polar night and subsequent successful return to light in the diatom *Fragilariopsis cylindrus*. New Phytologist 241, 2193–2208.

Jouhet J, Lupette J, Clerc O, Magneschi L, Bedhomme M, Collin S, Roy S, Maréchal E, Rébeillé F. 2017. LC-MS/MS versus TLC plus GC methods: Consistency of glycerolipid and fatty acid profiles in microalgae and higher plant cells and effect of a nitrogen starvation. (I-F Chang, Ed.). PLOS ONE 12, e0182423.

Lacour T, Morin P-I, Sciandra T, Donaher N, Campbell DA, Ferland J, Babin M. 2019. Decoupling light harvesting, electron transport and carbon fixation during prolonged darkness supports rapid recovery upon re-illumination in the Arctic diatom Chaetoceros neogracilis. Polar Biology 42, 1787–1799.

Leyland B, Novichkova E, Dolui AK, Jallet D, Daboussi F, Legeret B, Li Z, Li-Beisson Y, Boussiba S, Khozin-Goldberg I. 2024. Acyl-CoA binding protein is required for lipid droplet degradation in the diatom *Phaeodactylum tricornutum*. Plant Physiology 194, 958– 981.

Li W, Wang R, Li M, Li L, Wang C, Welti R, Wang X. 2008. Differential degradation of extraplastidic and plastidic lipids during freezing and post-freezing recovery in *Arabidopsis thaliana*. Journal of Biological Chemistry 283, 461–468.

Li Z, Zhang Y, Li W, Irwin AJ, Finkel ZV. 2023. Common environmental stress responses in a model marine diatom. New Phytologist 240, 272–284.

Mann KH, Lazier JRN. 2013. Dynamics of marine ecosystems: biological-physical interactions in the oceans. Malden, MA: Blackwell Pub.

Massana R, del Campo J, Sieracki ME, Audic S, Logares R. 2014. Exploring the uncultured microeukaryote majority in the oceans: reevaluation of ribogroups within stramenopiles. The ISME Journal 8, 854–866.

Matthijs M, Fabris M, Obata T, Foubert I, FrancoZorrilla JM, Solano R, Fernie AR, Vyverman W, Goossens A. 2017. The transcription factor bZIP14 regulates the TCA cycle in the diatom *Phaeodactylum tricornutum*. The EMBO Journal 36, 1559–1576.

Mellor GL, Durbin PA. 1975. The structure and dynamics of the ocean surface mixed layer. Journal of Physical Oceanography, 718–728.

Mills JK, Needham D. 2005. Lysolipid incorporation in dipalmitoylphosphatidylcholine bilayer membranes enhances the ion permeability and drug release rates at the membrane phase transition. Biochimica et Biophysica Acta (BBA) - Biomembranes 1716, 77–96.

Miquel M, James D, Dooner H, Browse J. 1993. *Arabidopsis* requires polyunsaturated lipids for low-temperature survival. Proceedings of the National Academy of Sciences 90, 6208–6212.

Moellering ER, Muthan B, Benning C. 2010. Freezing tolerance in plants requires lipid remodeling at the outer chloroplast membrane. Science 330, 226–228.

Morgan-Kiss RM, Priscu JC, Pocock T, Gudynaite-Savitch L, Huner NPA. 2006. Adaptation and acclimation of photosynthetic microorganisms to permanently cold environments. Microbiology and molecular biology reviews: MMBR 70, 222–252.

Morin P, Lacour T, Grondin P, et al. 2020. Response of the sea[ice diatom *Fragilariopsis cylindrus* to simulated polar night darkness and return to light. Limnology and Oceanography 65, 1041–1060.

Mortensen SH, Børsheim KY, Rainuzzo J, Knutsen G. 1988. Fatty acid and elemental composition of the marine diatom *Chaetoceros gracilis* Schütt. Effects of silicate deprivation, temperature and light intensity. Journal of Experimental Marine Biology and Ecology 122, 173–185.

Moustafa A, Beszteri B, Maier UG, Bowler C, Valentin K, Bhattacharya D. 2009. Genomic footprints of a cryptic plastid endosymbiosis in diatoms. Science 324, 1724–1726.

Murata N, Yamaya J. 1984. Temperature-dependent phase behavior of phosphatidylglycerols from chilling-sensitive and chilling-resistant plants. Plant Physiology 74, 1016–1024.

Nisbet RER, Kilian O, McFadden GI. 2004. Diatom genomics: Genetic acquisitions and mergers. Current Biology 14, R1048–R1050.

Nymark M, Valle KC, Hancke K, Winge P, Andresen K, Johnsen G, Bones AM, Brembu T. 2013. Molecular and Photosynthetic Responses to Prolonged Darkness and Subsequent Acclimation to Re-Illumination in the Diatom Phaeodactylum tricornutum. (R Subramanyam, Ed.). PLoS ONE 8, e58722.

Raven JA, Kübler JE, Beardall J. 2000. Put out the light, and then put out the light. Journal of the Marine Biological Association of the United Kingdom 80, 1–25.

Reagan JR, Boyer TP, García HE, et al. 2023. World Ocean Atlas 2023.

Rogato A, Amato A, Iudicone D, Chiurazzi M, Ferrante MI, d’Alcalà MR. 2015. The diatom molecular toolkit to handle nitrogen uptake. Marine Genomics 24, 95–108.

Routaboul J-M, Fischer SF, Browse J. 2000. Trienoic fatty acids are required to maintain chloroplast function at low temperatures. Plant Physiology 124, 1697–1705.

Seeleuthner Y, Mondy S, Lombard V, et al. 2018. Single-cell genomics of multiple uncultured stramenopiles reveals underestimated functional diversity across oceans. Nature Communications 9, 310.

Seiwert D, Witt H, Janshoff A, Paulsen H. 2017. The non-bilayer lipid MGDG stabilizes the major light-harvesting complex (LHCII) against unfolding. Scientific Reports 7, 5158.

Shimojima M, Ohta H. 2022. Plant and algal galactolipids: Their function, biosynthesis and evolution. Advances in Botanical Research. Elsevier, 59–89.

Shomo ZD, Li F, Smith CN, Edmonds SR, Roston RL. 2024. From sensing to acclimation: The role of membrane lipid remodeling in plant responses to low temperatures. Plant Physiology 196, 1737–1757.

Sklar LA, Miljanich GP, Dratz EA. 1979. Phospholipid lateral phase separation and the partition of cis-parinaric acid and trans-parinaric acid among aqueous, solid lipid, and fluid lipid phases. Biochemistry 18, 1707–1716.

Sorhannus U. 2007. A nuclear-encoded small-subunit ribosomal RNA timescale for diatom evolution. Marine Micropaleontology 65, 1–12.

Steponkus PL. 1984. Role of the plasma membrane in freezing injury and cold acclimation. Annual Review of Plant Physiology 35, 543–584.

Svenning JB, Vasskog T, Campbell K, Bæverud AH, Myhre TN, Dalheim L, Forgereau ZL, Osanen JE, Hansen EH, Bernstein HC. 2024. Lipidome plasticity enables unusual photosynthetic flexibility in Arctic vs. temperate diatoms. Marine Drugs 22, 67.

Sverdrup HU. 1953. On conditions for the vernal blooming of phytoplankton. ICES Journal of Marine Science 18, 287–295.

Thaler M, Lovejoy C. 2014. Environmental selection of marine stramenopile clades in the Arctic Ocean and coastal waters. Polar Biology 37, 347–357.

Thompson PA, Guo M, Harrison PJ. 1992a. Effects of variation in temperature. I. on the biochemical composition of eight species of marine phytoplankton. Journal of Phycology 28, 481–488.

Thompson PA, Guo M, Harrison PJ, Whyte JNC. 1992b. Effects of variation in temperature. II. on the fatty acid composition of eight species of marine phytoplankton. Journal of Phycology 28, 488–497.

Tiku PE, Gracey AY, Macartney AI, Beynon RJ, Cossins AR. 1996. Cold-induced expression of Δ^9^-Desaturase in carp by transcriptional and posttranslational mechanisms. Science 271, 815–818.

Villar E, Zweig N, Vincens P, et al. 2025. DIATOMICBASE[: a versatile gene[centered platform for mining functional omics data in diatom research. The Plant Journal 121, e70061.

Wang J, Liu Z, Liu H, Peng D, Zhang J, Chen M. 2021. *Linum usitatissimum* FAD2A and FAD3A enhance seed polyunsaturated fatty acid accumulation and seedling cold tolerance in *Arabidopsis thaliana*. Plant Science 311, 111014.

Willette S, Gill SS, Dungan B, Schaub TM, Jarvis JM, St. Hilaire R, Omar Holguin F. 2018. Alterations in lipidome and metabolome profiles of *Nannochloropsis salina* in response to reduced culture temperature during sinusoidal temperature and light. Algal Research 32, 79–92.

Wu J, Nadeem M, Galagedara L, Thomas R, Cheema M. 2022. Recent insights into cell responses to cold stress in plants: Signaling, defence, and potential functions of phosphatidic acid. Environmental and Experimental Botany 203, 105068.

Zhang H, Xiong X, Guo K, et al. 2024. A rapid aureochrome opto-switch enables diatom acclimation to dynamic light. Nature Communications 15, 5578.

Zheng G, Li L, Li W. 2016. Glycerolipidome responses to freezing- and chilling-induced injuries: examples in Arabidopsis and rice. BMC Plant Biology 16, 70.

Zheng G, Tian B, Zhang F, Tao F, Li W. 2011. Plant adaptation to frequent alterations between high and low temperatures: remodelling of membrane lipids and maintenance of unsaturation levels. Plant, Cell & Environment 34, 1431–1442.

Zhu J, Li S, Chen W, et al. 2024. Delta-5 elongase knockout reduces docosahexaenoic acid and lipid synthesis and increases heat sensitivity in a diatom. Plant Physiology 196, 1356– 1373.

Zorin B, Pal-Nath D, Lukyanov A, Smolskaya S, Kolusheva S, Didi-Cohen S, Boussiba S, Cohen Z, Khozin-Goldberg I, Solovchenko A. 2017. Arachidonic acid is important for efficient use of light by the microalga *Lobosphaera incisa* under chilling stress. Biochimica et Biophysica Acta (BBA) - Molecular and Cell Biology of Lipids 1862, 853–868.

